# Auditory predictions and prediction errors in response to self-initiated vowels

**DOI:** 10.1101/671990

**Authors:** Franziska Knolle, Michael Schwartze, Erich Schröger, Sonja A. Kotz

## Abstract

It has been suggested that speech production is accomplished by an internal forward model, reducing processing activity directed to self-produced speech in the auditory cortex. The current study uses an established N1-suppression paradigm comparing self- and externally-initiated natural speech sounds to answer two questions:

1. Are forward predictions generated to process complex speech sounds, such as vowels, initiated via a button press?
2. Are prediction errors regarding self-initiated deviant vowels reflected in the corresponding ERP components?

Results confirm an N1-suppression in response to self-initiated speech sounds. Furthermore, our results suggest that predictions leading to the N1-suppression effect are specific, as self-initiated deviant vowels do not elicit an N1-suppression effect. Rather, self-initiated deviant vowels elicit an enhanced N2b and P3a compared to externally-generated deviants, externally-generated standard, or self-initiated standards, again confirming prediction specificity.

Results show that prediction errors are salient in self-initiated auditory speech sounds, which may lead to more efficient error correction in speech production.

## Introduction

Speaking is a highly complex human capacity: It does not only involve a motor act, but also leads to the perception and the monitoring of one’s own voice. The distinction between self-produced speech from speech of others is proposed to be accomplished by a “motor-to-sensory discharge” (Paus, Perry, Zatorre, Worselyy, & Evans, 1996) or an internal forward model (Hickok, 2012; Tian & Poeppel, 2010; Ventura, Nagarajan, & Houde, 2009). The idea of an internal forward model suggests that an efference copy (von Holst & Mittelstädt, 1950) of a motor act is generated that predicts its sensory consequences (Wolpert, Ghahramani, & Jordan, 1995). The prediction prepares a respective cortical area to perceive the predicted sensory input. Consequently, brain activity directed to incoming sensation is suppressed (Chen et al, 2011).

Interestingly, the suppression effect has been reported in many vocalization studies. It was shown that speech production elicits smaller event-related potentials (ERPs) or fields (ERFs) than passively perceived speech (Curio, Neuloh, Numminen, Jousmaki, & Hari, 2000; Ford, Gray, Faustman, Roach, & Mathalon, 2007; Gunji, Hoshiyama, & Kakigi, 2001; Numminen & Curio, 1999; Numminen, Salmelin, & Hari, 1999; Ott & Jäncke, 2013; Ventura et al., 2009). Based on a non-human primate study (Müller-Preuss & Ploog, 1981; replication and extension of non-human primate investigations by Eliades & Wang, 2003), Creutzfeldt and colleagues (1989) recorded intracranial neuronal activity from the right and left superior, middle and inferior temporal gyri in patients undergoing surgery for epilepsy. Results revealed suppressed activity in response to vocalization. Relatedly, Chen and colleagues (2011) conducted an electrocorticography (ECoG) study during human vocalization. They reported neural phase synchrony in the gamma band between Broca’s area and the auditory cortex. This phase synchrony that preceded a speaker’s speech onset was greater during vocalizing than when listening to their own speech passively (i.e., pre-recorded), indicating that phase synchrony in the gamma band between the two brain regions may describe the transmission of a motor efference copy.

In a PET-study, Hirano and colleagues (1996, 1997) found strong cerebellar activation when investigating self-produced speech that was either normal or distorted (i.e., delayed or changed in pitch), possibly indicating the role of the cerebellum in generating internal forward predictions based on an efference copy. Along similar lines, we have shown that the cerebellum is involved in generating motor-to-auditory predictions when processing self-initiated sounds (Knolle, Schröger, Baess, & Kotz, 2012; Knolle, Schröger, & Kotz, 2013a). We utilized a N1-suppression paradigm (Schäfer & Marcus, 1973) to compare self-initiated (via finger tap) with externally-generated sinusoidal tones. We found that patients with focal cerebellar lesions did not show a significant N1-suppression effect in response to self-initiated sounds. This indicates that the cerebellum is involved in generating auditory forward predictions – or their (sensory) attenuation.

In a further study (Knolle, Schröger & Kotz, 2013b), we compared self- and externally-generated sounds including 30% unexpected deviant, or ‘oddball’, sounds (i.e., sounds altered in frequency). We investigated the violation of a prediction when processing a self-generated deviant sound; furthermore we investigated the reflection of such prediction errors in the ERP. The results revealed that precise predictions concerning the acoustic features of a self-generated sound were formed (i.e., modulated N1 suppression). The result was supported by an enhancement of “auditory responsiveness” towards a self-generated deviant shown in an increase in amplitude of the N2b and P3a components. The increased saliency of self-generated deviant sounds may lead to more efficient processing compared to externally-generated deviants. In other words, the increased salience of certain stimuli may reverse the sensory attenuation induced by agency. Thus, the study provides a more complete picture concerning auditory forward prediction and prediction errors with respect to self-generated sinusoidal sounds. Note that the detection of prediction errors is not in the service of detecting the content or cause of a stimulus – but in evaluating how predictable that stimulus was. This is a subtle but important point, which speaks to the predictive processing of the precision or predictability of different stimuli in different settings.

Based on this study the question arose, if precise predictions are also generated in response to complex auditory stimuli, such as speech sounds. Former studies (Behroozmand, Liu, & Larson, 2011; Behroozmend, Sangtina, Korzyukov, & Larson, 2016; Christoffels, van de Ven, Formisano, & Schiller, 2011; Fu et al., 2006; Heinks-Maldonado, Mathalon, Gray, & Ford, 2005; Heinks-Maldonado, Nagarajan, & Houde, 2006) which compared natural and altered self-produced vocalizations suggest that this is indeed the case. They reported a reduced N1-suppression effect in response to altered compared to unchanged vocal feedback, indicating the generation of a precise prediction. Although altered auditory feedback creates prediction errors, these studies did not aim to investigate the detection of deviance (i.e., altered auditory feedback), nor have they discussed feedback alterations in terms of prediction errors. Although these studies used a carefully conducted design which aimed at reducing as many confounding factors as possible (i.e., participants were asked to hold head, jaw, and tongue in a stationary position to reduce muscle contraction to a minimum), an inherent limitation of these studies is that it is impossible to control for motor activity induced by self-produced vocalization. Hence, a motor command is conducted in order to produce the sound. Based on the motor command, a sensory input is predicted. Consequently, the motor command may influence the ERP in response to the auditory output. Thus, we believe that an appropriate motor control condition (e.g., internal sound production) is necessary in order to control for such effects in the auditory ERPs. Furthermore, Heinks-Maldonado et al. (2005) point out that the differences in sound quality between a speaking and a listening condition could substantially influence the results, and possibly create the suppression effect. Nonetheless, these findings provide support the notion that predictions are generated based on precise patterns, also capturing complex auditory stimuli. However, as the motor-to-auditory links are much tighter with self-vocalization than the motor-to-auditory links involved when generating a sound via a finger tap, it is of interest whether the results obtained with self-vocalization can be generalized to other forms of self-generation of speech sounds.

In contrast to previous studies, the current study investigated whether predictions are formed on a precise patterns also in response to manually initiated complex, natural sounds (i.e., speech sounds), and whether the generation of a precise prediction leads to differential processing of self-compared to externally-produced complex deviant stimuli, seen in prediction errors specific to stimuli type. In a first experiment, a standard N1-suppression paradigm was used which has been studied extensively using click sounds, sinusoidal sounds or complex instrumental sounds (Baess, Jacobson, & Schröger, 2008; Baess, Horváth, Jacobson, & Schröger, 2011; Knolle et al., 2012; Lange, 2011; Martikainen, Kaneko, & Hari, 2005 (MEG); McCarthy & Donchin, 1976; Schäfer & Marcus, 1973). Here, we compared complex, natural vowels that were self-initiated via a finger tap to the same vowels, externally-produced (i.e., pre-recorded vowels /a:/), similarly to a recent study by Pinheiro and colleagues (Pinheiro, Schwartze & Kotz, 2018). This first experiment examined whether the N1-suppression paradigm was applicable to complex stimuli, such as speech sounds in a highly controlled setup. Based on our preceding studies (Knolle et al., 2012; Knolle et al., 2013a) investigating sinusoidal tones in a N1-suppression paradigm, we expected to find a N1-suppression followed by a P2-reduction in response to self-initiated vowels. Whereas the N1-suppression may reflect the unconscious, automatic formation of a prediction, preparing the auditory cortex to receive sensory input, the P2-reduction may reveal a later, more conscious processing stage of the generation of a prediction (i.e., the conscious detection of a self-initiated vowel).

In a second experiment, we investigated the violation of a precise prediction regarding self-generated complex speech sounds by introducing surprising, unpredictable events, and furthermore we explored the reflection of such prediction errors in the corresponding ERP components. Thus, we modified the N1-suppression paradigm of experiment one corresponding to a former study on self- and externally-generated deviant sounds (Knolle et al., 2013b). We compared self-initiated and externally-produced natural vowels, of which 30% were altered in quality (either /a:/ which is an open front unrounded vowel, or /o:/ which is a mid-close back rounded vowel), creating self-initiated and externally-produced deviants (Knolle et al., 2013b). If precise predictions were generated to process self-initiated vowels, the N1-suppression effect should be modified when a self-initiated deviant vowel is elicited. Furthermore, if self-initiated deviant vowels which created a prediction error were more salient than externally-produced deviant vowels they should reveal an enhanced N2b and P3a (Knolle et al., 2013b). These results would provide additional support for the generation of a precise prediction, which impacts the neural suppression and the detection of prediction errors in self-generated complex speech sounds.

## Methods

### Participants

Sixteen volunteers (8 females) participated in the current study. All participants were right-handed according to the Edinburgh Handedness Inventory (Oldfield, 1971). The mean age was 24.9 years (SD: 1.8 years) and ranged from 23 to 27 years. Participants were students of the University of Leipzig and were recruited via the participants’ database of the Max-Planck Institute for Human Cognitive and Brain Sciences, Leipzig, Germany. None of the participants reported any neurological dysfunction, but normal or corrected-to-normal visual acuity, and normal hearing. All participants gave their written informed consent and were paid for their participation. The study was conducted in accordance with the Declaration of Helsinki and approved by the Ethics Committee of the Leipzig University.

### Speech Stimuli

In order to obtain individual vowels, we recorded vowel samples from all participants using the program AlgoRec™ TerraTec Edition. Thus, throughout the experiment each participant listened to the vowels that they had previously produced. We asked the participants to produce “ah” and “oh” as in the German words /a:b<u>ɐ</u>/ (aber; engl. but) and /o:b<u>ɐ</u>/ (Ober; engl. waiter). We recorded 10 trials for each vowel per participant. For each participant we picked the vowel that best matched the average characteristics within the 10 trials. Using Praat 5.2.03 (1992-2010 by Paul Boersma and David Weenink; University of Amsterdam, Amsterdam, NL) we applied minimal normalization procedures to duration, intensity and pitch to maintain natural sound quality. The vowel duration of /a:/ was approximately 360.69ms (SD: 65.66ms) and pitch around 77.44 Hz (SD: 3.80 Hz). The vowel /o:/ had an average duration of 366.31ms (SD: 77.75ms) and an average pitch of 80.76 Hz (SD: 3.84 Hz). For both vowels, the sound intensity was calibrated at about 80 dB SPL and an average loudness of 70 dB was maintained by all subjects.

### Experimental conditions – Experiment 1

The first experiment contained two experimental conditions and one control condition (Figure 1, white background). In the vowel-motor condition (VMC-1) participants induced finger taps about every 2.4s (see Knolle et al., 2012 for a detailed description of the paradigm). Each tap elicited an immediate presentation of the vowel /a:/ (delay of 2-4ms due to the loading of the stimuli) via headphones. The acoustic stimulation, including self-initiated vowels, was recorded online and used as an ’external vowel sequence’ in the vowel-only condition (VOC-1). During VOC-1 participants did not produce finger taps, but were simply asked to listen and attend to the vowels. Lastly, participants carried out a motor-only condition (MOC-1), in which they also performed self-paced finger taps every 2.4s. However, in contrast to VMC-1, no sound was induced via the finger tap. This condition controlled for motor activity in VMC-1.

**Figure 1.**
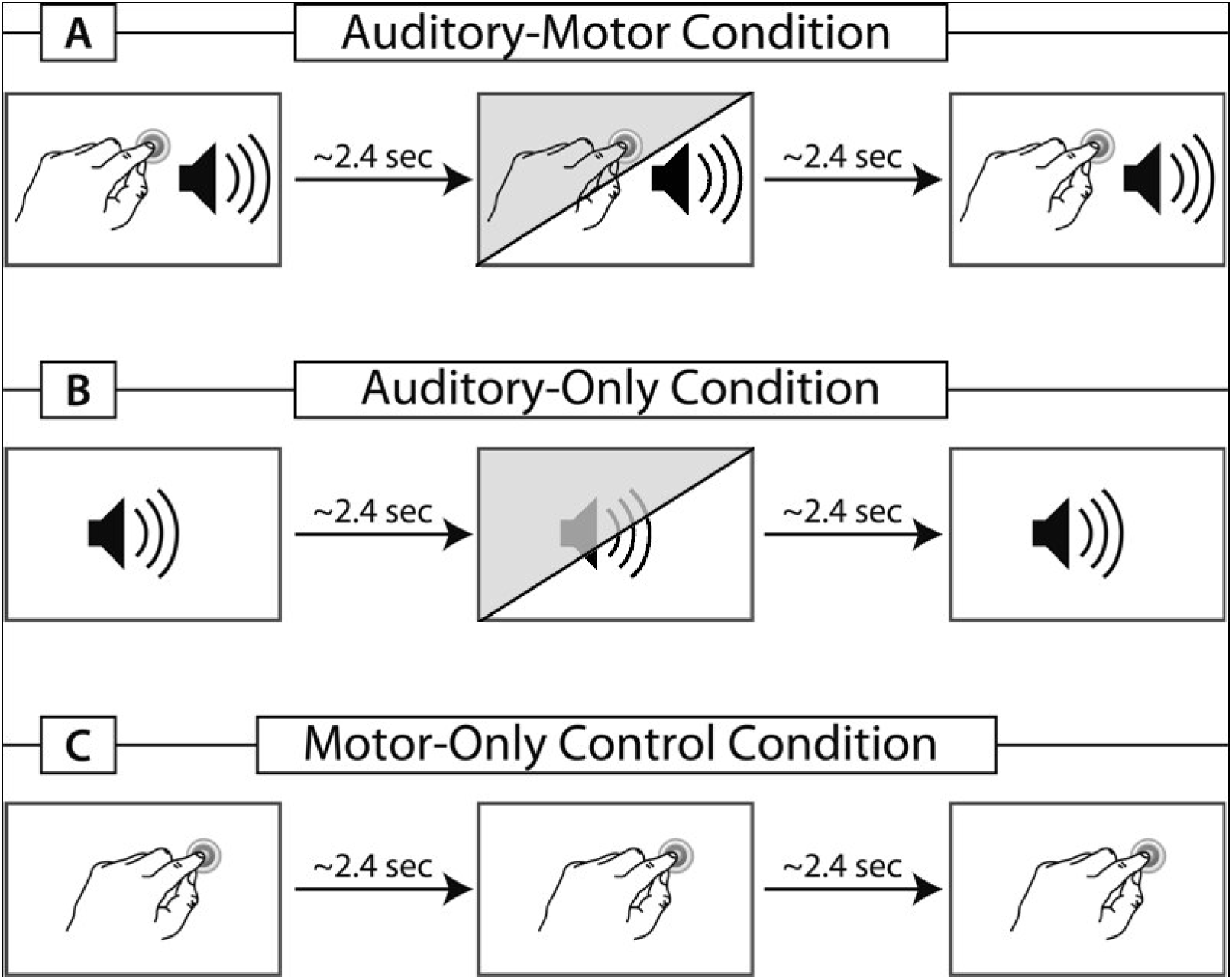
Schematic illustration of the three different conditions used in the studies. A) represents the vowel-motor condition (VMC-1/2): A vowel is self-initiated via a finger tap. In experiment 2, 30% of the finger taps elicit a vowel deviant, indicated in the illustration via the diagonal shading. B) represents the vowel-only condition (VOC-1/2): The vowel sequence is presented externally, containing all vowels from the VMC-1/2 accordingly. The diagonal shading presents the inclusion of deviant vowels used in experiment 2. C) illustrates the motor-only control condition (MOC-1/2): taps are required, but no sound is elicited. The motor-only condition is identical in experiment 1 and 2.

### Experimental conditions – Experiment 2

In experiment 2, two experimental and one control condition were presented (Figure 1b). In the vowel-motor condition (VMC-2) participants induced finger taps about every 2.4s. Each tap elicited an immediate presentation of either the vowel /a:/ or /o:/ via headphones. In 30% of the taps a deviant was presented. If the standard stimulus was the vowel /a:/, the deviant was the vowel /o:/ and vice versa. The timing of the acoustic stimulation was recorded online. This information was used to produce an identical but externally generated vowel sequence in the vowel-only condition (VOC-2). Thus, participants received exactly the same set of stimuli in both experimental conditions. During VOC-2 participants did not produce finger taps, but were simply asked to listen attentively to the auditory stimuli. As in experiment 1 MOC-2 served as a control condition for motor activity in VMC-2.

Both experimental runs were preceded by two training blocks each. In the first block, participants practiced to tap every 2.4s. The second training block included visual feedback to indicate whether a trial was too slow (tapping interval longer than 3s) or too fast (tapping interval shorter than 1.8s). The feedback ensured that participants had learned to estimate the time between two successive finger taps without counting. Trials outside the range of 1.8s – 2.4s were treated as errors. During the experimental run, no feedback was given.

### Experimental procedure

Participants were comfortably seated in an electrically shielded and sound-attenuated experimental chamber. A fixation cross was displayed in the middle of a computer screen. To ensure that the motor activity was comparable across participants in the auditory-motor and the motor-only condition, they were instructed to change hands (index finger) whenever indicated on the screen. Hence all participants tapped in equal parts with left and right hand. The order of tapping hands was randomized across participants. Each tap triggered the instantaneous presentation of a vowel via headphones (Sennheiser HD 202) to both ears in VMC-1/2 and VOC-1/2. An in-house built, highly sensitive tapping device was used to record the finger taps. Participants wore headphones in order to cover them up from all sounds possibly emitted by the taps. In the second experiment, the participants performed a combination of VMC-2 and VOC-2: during one run the standard vowel that was triggered via a tap was the vowel /a:/ and the deviant was the vowel /o:/. In the other run, the allocation of standard and deviant vowel was a reversed. The first experiment consisted of 100 trials in each condition. Additionally, we collected 100 trials in MOC-1/2. In the second experiment, we recorded 200 trials in each the VMC-2 and VOC-2 with 70% /a:/ as standard and 30% /o:/ as deviant and vice versa. In total, 700 trials were recorded in both experiments. Experimental conditions were presented in blocks of 100 trials each. Block order was restricted: The VMC-1/2 always preceded VOC-1/2, but the MOC-1/2 was randomized across participants.

### Electrophysiological recordings

The electroencephalogram (EEG) was recorded continuously from 59 Ag–AgCl electrodes according to the International 10–20 system. In addition, activity from the left and right mastoids and the sternum (ground electrode) was recorded. The EEG was sampled at a rate of 500Hz (Refa amplifiers system, TMS international, Enschede, NL) and an anti-aliasing filter of 135 Hz was applied. To control for eye movements, vertical and horizontal electrooculograms (EOG) were recorded bipolarly. The impedance of all electrodes was kept below 5 kΩ. The recordings were online referenced to the left mastoid. EEP 3.2.1 Max-Planck-Institute of Cognitive Neuroscience, Leipzig, Germany was used to process the data.

### Data Analysis – Behavioral Data

Tapping intervals shorter than 1.8s or longer than 3.0s were treated as errors, and were excluded from further EEG analysis. We acquired tapping intervals for VMC-1/2 and MOC-1/2 using the Presentation software (Neurobehavioral Systems, Inc., Albany, CA, USA). For each participant we generated the mean length of the tapped interval per condition and the overall performance accuracy (percent correct; ACCURACY) separately for VMC-1/2 and MOC-1/2.

### Data Analysis – EEG Data

The EEG data were filtered with a 0.3-15Hz bandpass filter (1601 Hamming windowed filter). The EEG data were re-referenced to linked mastoids. ERPs were time-locked to the stimulus onset of all critical trials. Each analyzed epoch lasted 600 ms including a 100 ms pre-stimulus baseline. The critical epochs were automatically scanned to reject horizontal and vertical eye-movements, muscle artifacts, and electrode drifts. Trials exceeding 30 µV at the eye channels and 40 µV at CZ were rejected. This automatic rejection was corrected manually by applying an eye-movement correction.

We controlled for motor activity by computing a difference-wave between VMC-1/2 and MOC-1/2 to compare the sensory activity elicited in the two experimental conditions. This corrected condition was labeled vowel-corrected condition (VCC-1, VCC-2). In the first experiment, we only compared fully predictable self-initiated and externally-produced vowels (i.e., /a:/). In the second experiment, we investigated two types of self-initiated vowels – standard (VCS-2) and deviant (VCD-2) vowels, as well as two types of externally-produced vowels – self-initiated standard and deviant vowels (VOS-2; VOD-2). As the statistical analysis of the ERP-results in response to standard vowel /a:/ and standard vowel /o:/ as well as to deviant vowel /a:/ and deviant vowel /o:/ did not differ significantly, we combined the standard vowels /a:/ and /o:/ as well as deviant vowel /a:/ and /o:/ for all further statistical analyses.

Group-average ERPs were generated for standards and deviants. In the first experiment, the difference waves (i.e., responses to externally generated minus self-initiated vowels) revealed two ERP responses, a negative one in the time window of the N1 peaking at approximately 90ms, and a positive response in the P2 time window peaking at approx. 190ms. Statistical analyses were calculated based on individual amplitudes, in the time windows of 70-110ms for the N1 and 170-210 ms for the P2. In experiment 2, difference waves revealed two ERP responses for standard vowels: a negative one in the N1 time window, peaking at approx. 90 ms followed by a positive one in the P2 time window peaking at approx. 190ms; and four ERP responses to deviant vowels: a N1 and P2, as well as an N2b peaking at approx. 170 ms and a later positive response in the time window of the P3a peaking at approx. 310ms. Statistical analyses were calculated based on individual amplitudes in the following time windows: 70-110 ms for the N1, 150-190 ms for the N2b, 170-210 ms for the P2, and 260-360 ms for the P3a. Furthermore, we applied a regions of interest (ROI) analysis using five ROIs (central (ZZ): FZ, CZ, PZ, FCZ, CPZ; left lateral (LL): F7, T7, P7, FT7, TP7; left medial (LM): F3, C3, P3, FC3, CP3; right lateral (RL): F8, T8, P8, FT8, TP8; right medial (RM): F4, C4, P4, FC4, CP4).

### Statistical analyses

For the statistical analysis, the SAS 8.20.20 (Statistical Analysis System, SAS Institute Inc., Cary, North Carolina, USA) software package was used. Only significant results are presented and where required, the Greenhouse–Geisser correction was applied. Furthermore, we conducted Bonferroni corrected post-hoc tests.

In the first experiment, comparing self-initiated /a:/-vowels to externally-produced /a:/-vowels, we ran two analyses of variance (ANOVA), one for each ERP component, including the within-subject factors CONDITION (self-initiated: VCC-1; externally-produced: VOC-1) and ROI (RL,RM, ZZ, LM, LL). For the statistical analyses of each individual ERP component in the second experiment, we ran a 2×2×5 ANOVA using the within-subject factors CONDITION (self-initiated: VCC-2; externally-produced: VOC-2), TYPE (standard vs. deviant) and ROI (LL, LM, RM, RL, ZZ).

## Results

### Behavioral data

In the first experiment, the average tapping interval duration was 2469.79 ms (SD: 360.77 ms) in VMC-1. Furthermore, participants tapped with an overall accuracy of 85.80% (SD: 18.73%). The Kolmogorov-Smirnov-Test revealed no deviance from the normal distribution (*p* = .18). In the second experiment, the results were similar. The average tapping interval was 2233.15 ms (SD: 257.43 ms) in VMC-2. Furthermore, participants tapped with an overall correctness of 92.92% (SD: 6.06%). The Kolmogorov-Smirnov-Test revealed a normal distribution (*p* = .82). In MOC-1/2 we found similar results, the average length of the tapping interval was 2388.94 ms (SD: 267.44 ms). Participants performed with an overall accuracy of 93.25% (SD: 8.14%; normal distribution: *p* = .52).

### ERP data – experiment 1

The statistical results are presented in Table 1, and summarized below. The CONDITION (self-initiated, externally-initiated) x ROI (LL, LM, ZZ, RM, RR) ANOVA revealed a significant effects in the time window of N1 and P2. In the N1 time window, results confirmed significant differences between the two conditions of self-initiated and externally initiated vowels; as well as significant interaction between conditions and region. Self-initiated vowels elicited a N1-suppression compared to externally-produced vowels (N1: mean amplitude VCC-1: −2.20 µV, mean amplitude VOC-1: −3.09 µV) (Figure 2).

**Table 1.** Results of Omnibus ANOVA in Experiment 1

| N1 |  |  | P2 |  |  |
| --- | --- | --- | --- | --- | --- |
|  | F | p |  | F | p |
| Condition<sup>a</sup> | 4.66 | .048 | Condition<sup>a</sup> | 10.16 | .006 |
| Condition x ROI<sup>b</sup> | 6.70 | <.001 | Condition x ROI<sup>b</sup> | 5.25 | .001 |

<sup>a</sup>F(1,15); <sup>b</sup>F(4,60)

**Figure 2.**
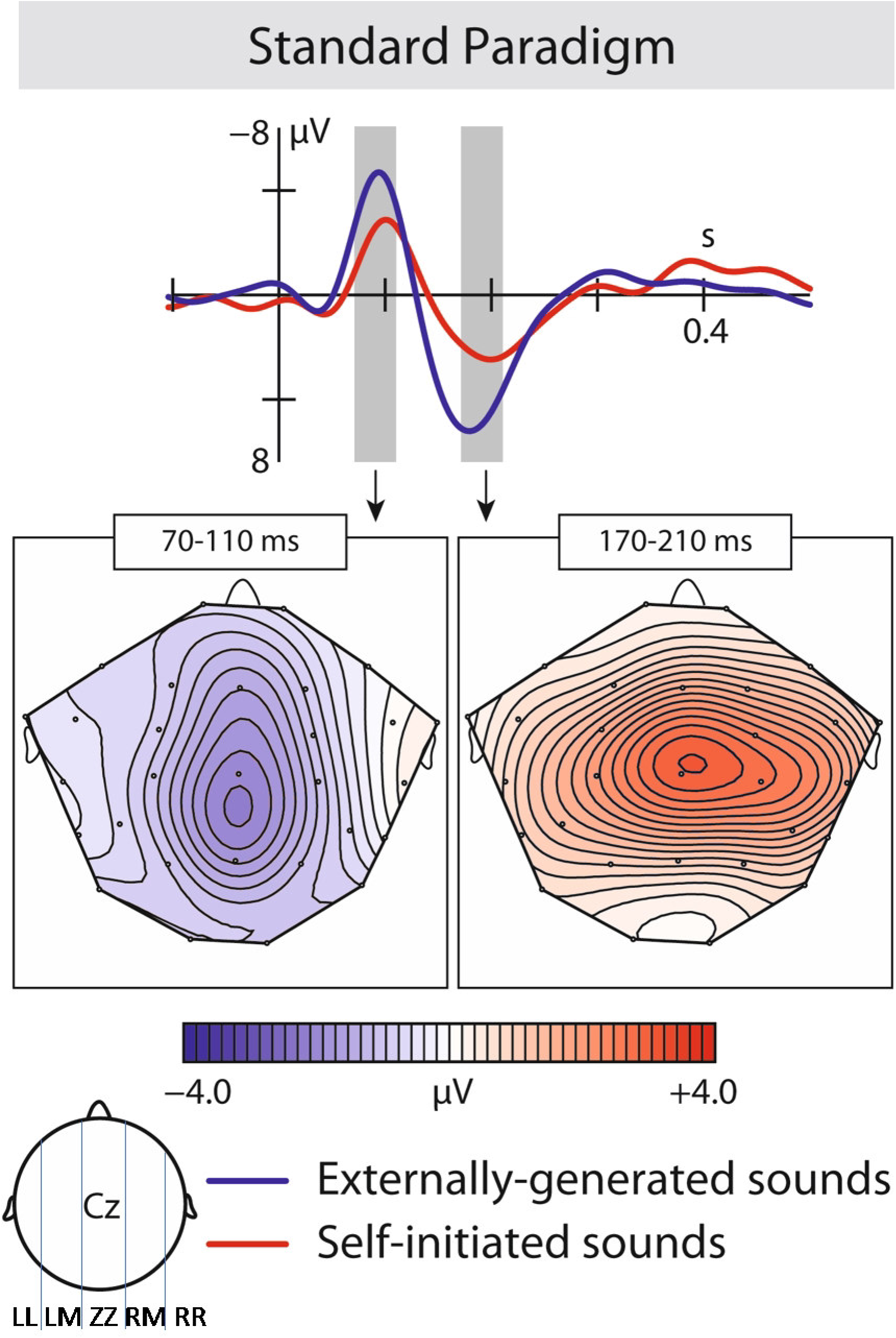
Results of the first experiment: ERP responses: Brain responses elicited by self-initiated and externally-produced vowels in the central region. The blue solid line represents externally-produced vowels (VOC-1), whereas the red solid line shows responses elicited by self-initiated vowels (VCC-1). Brain maps: Grand average scalp maps showing the spatial distribution of the difference waves (VOC-1 – VCC-1) in the analyzed N1 and P2 time window.

In the P2 time window, similarly to the N1, we found a significant difference between the two conditions and a significant interaction of condition and region, with self-initiated vowels being significantly suppressed compared to externally-produced vowels (P2: mean amplitude VCC-1: 1.73 µV, mean amplitude VOC-1: 3.44 µV) (Figure 2).

Experiment 1 revealed the expected N1-suppression effect as well as a reduced P2 in response to self-initiated vowels.

### ERP data – experiment 2

All statistical results are presented in Table 2. For all ERP components we conducted a CONDITION (self-initiated, externally-initiated) x ROI (LL, LM, ZZ, RM, RR) x TYPE (standard, deviant).

**Table 2.**
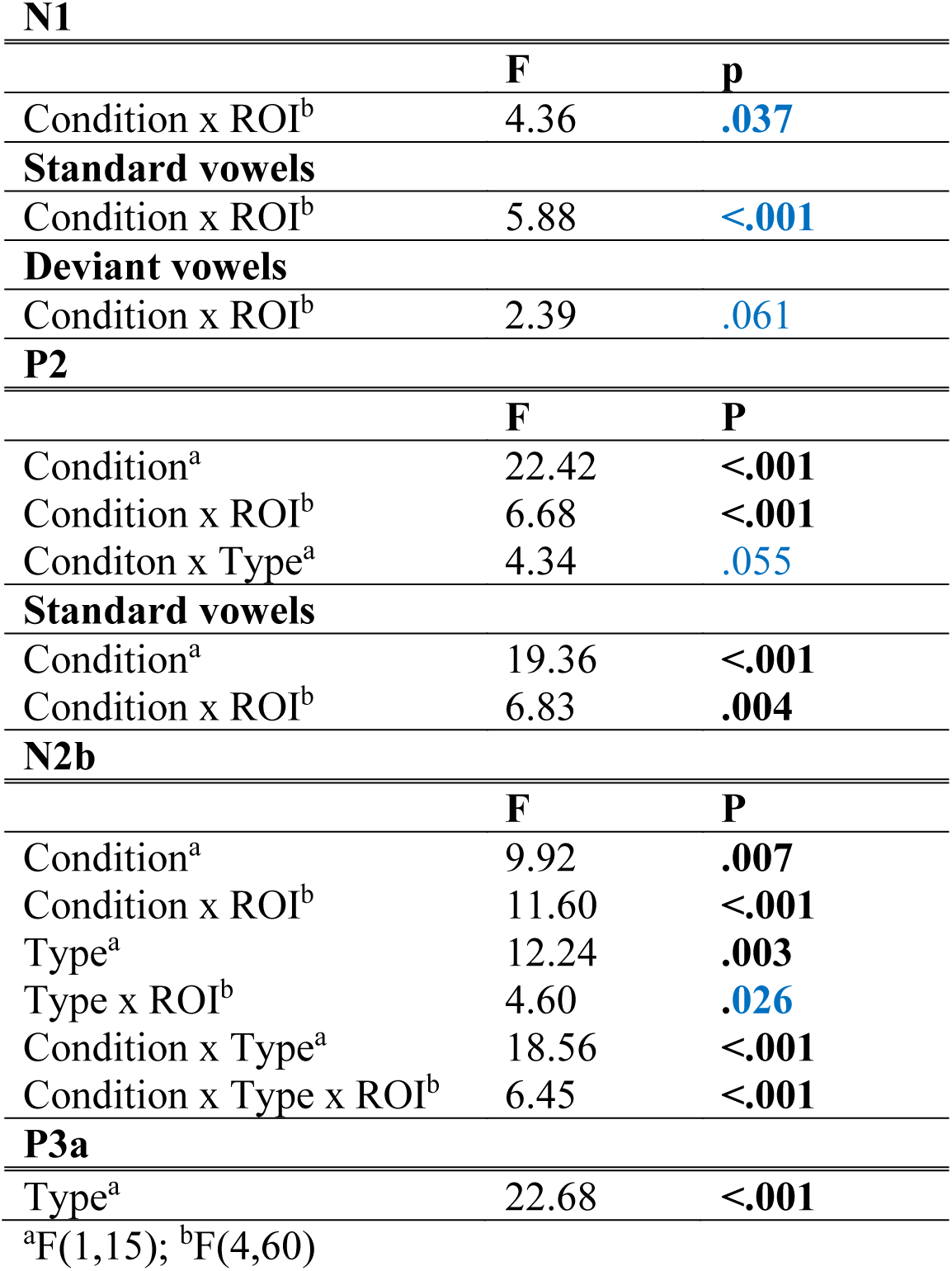
Results of Omnibus ANOVA in Experiment 2

In the N1 time window, the ANOVA revealed significant differences between condition and region. According to our hypothesis, we conducted a planned analysis, investigating condition effects within the different vowel types: In the standard vowels (Figure3a) we found a significant suppression effect in response to self-initiated vowels which was shown by an interaction of condition and region (mean amplitudes standard vowels: VCS-2 mean amplitude: −2.98 µV, VOS-2 mean amplitude: −3.77 µV). In the deviant vowels (Figure 3B), the suppression effect did not reach significance (mean amplitudes deviant vowels: VCD-2 mean amplitude: −3.81 µV, VOD-2 mean amplitude: −4.04 µV).

**Figure 3.**
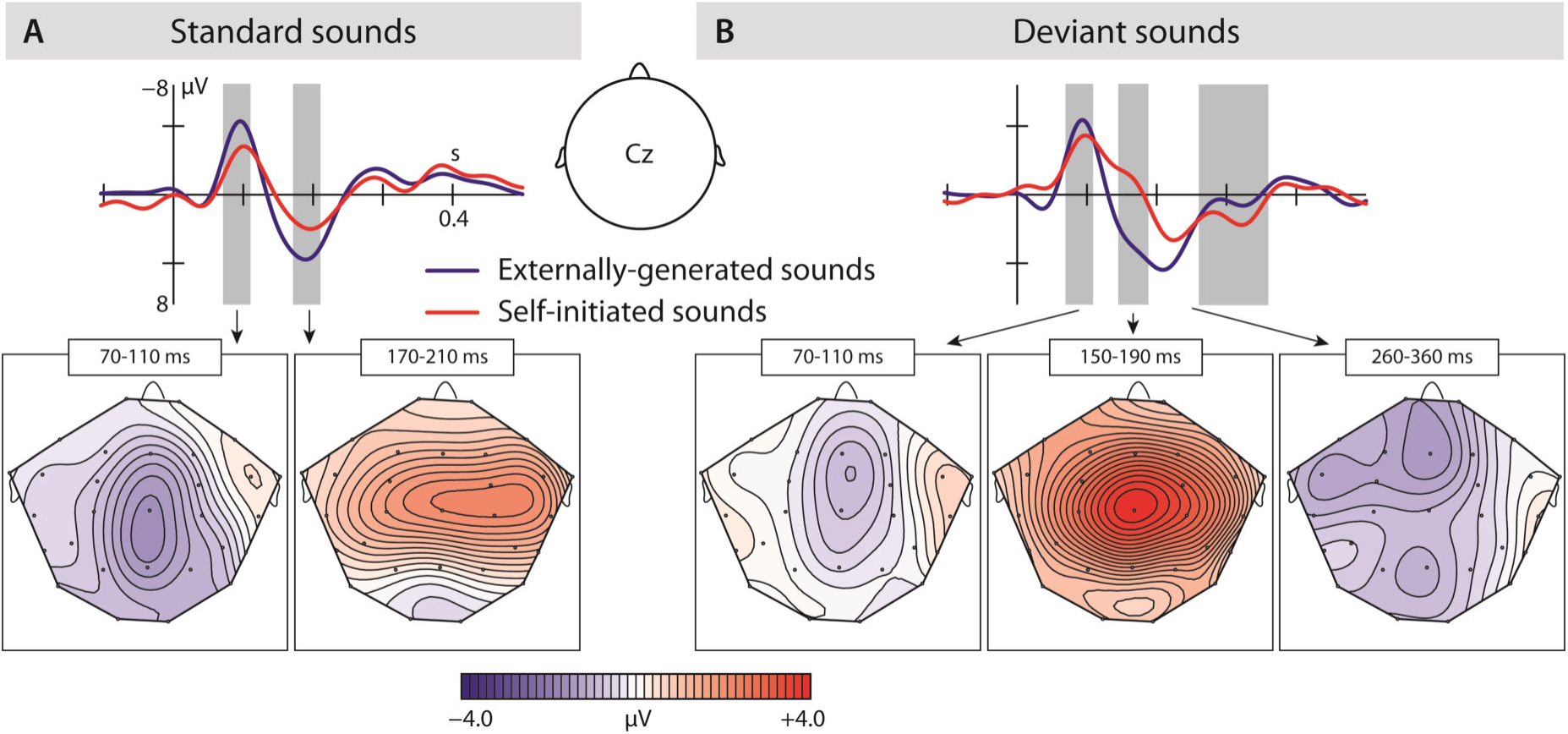
Results of the second experiment: A) Brain responses elicited by self-initiated and externally-produced standard vowels in the central region. The blue solid line represents externally-produced standard vowels (VOS-2), whereas the red solid line shows responses elicited by self-initiated standard vowels (VCS-2). B) Brain responses elicited by self-initiated and externally-produced deviant vowels also in the central region. The blue solid line shows externally-produced standard vowels (VOD-2), and the red solid line shows responses to self-initiated deviant vowels (VCD-2). The brain maps show the distribution of the effect (for standard vowels: VOS-2 minus VCS-2; for deviant vowels: VOD-2 minus VCD-2). Grey bar reflect analyzed in the time window of the N1, N2b and P3a.

The results show that we only find a N1-suppression effect in response to self-initiated standard.

In the P2 time window, the ANOVA analysis revealed a significant difference between conditions – self-initiated versus externally initiated – and types – standard versus deviants, as well as significant interactions between condition and region aa well as a marginally significant effect between condition and type. In response to standard vowels (Figure3a), we found a significant suppression effect in response to self-initiated vowels compared to externally initiated vowels (Standard: VCS-2 mean amplitude: 1.05 µV, VOS-2 mean amplitude: 2.62 µV) shown by a condition effect as well as a condition by region interaction. Deviant vowels showed a different pattern (Figure3b). In response to externally-generated deviant vowels we found a P2 (VOD-2 mean amplitude: 2.55 µV) that did not differ significantly from externally-produced standards (VOS-2 mean amplitude: 2.62 µV) (Figure 4A). The visual inspection of the ERPs in response to the self-initiated deviants showed a very small shoulder following the N1, which may reflect a P2. However, this component was overlaid by a strong N2b effect, which was elicited in the time window of the P2, and described below.

**Figure 4.**
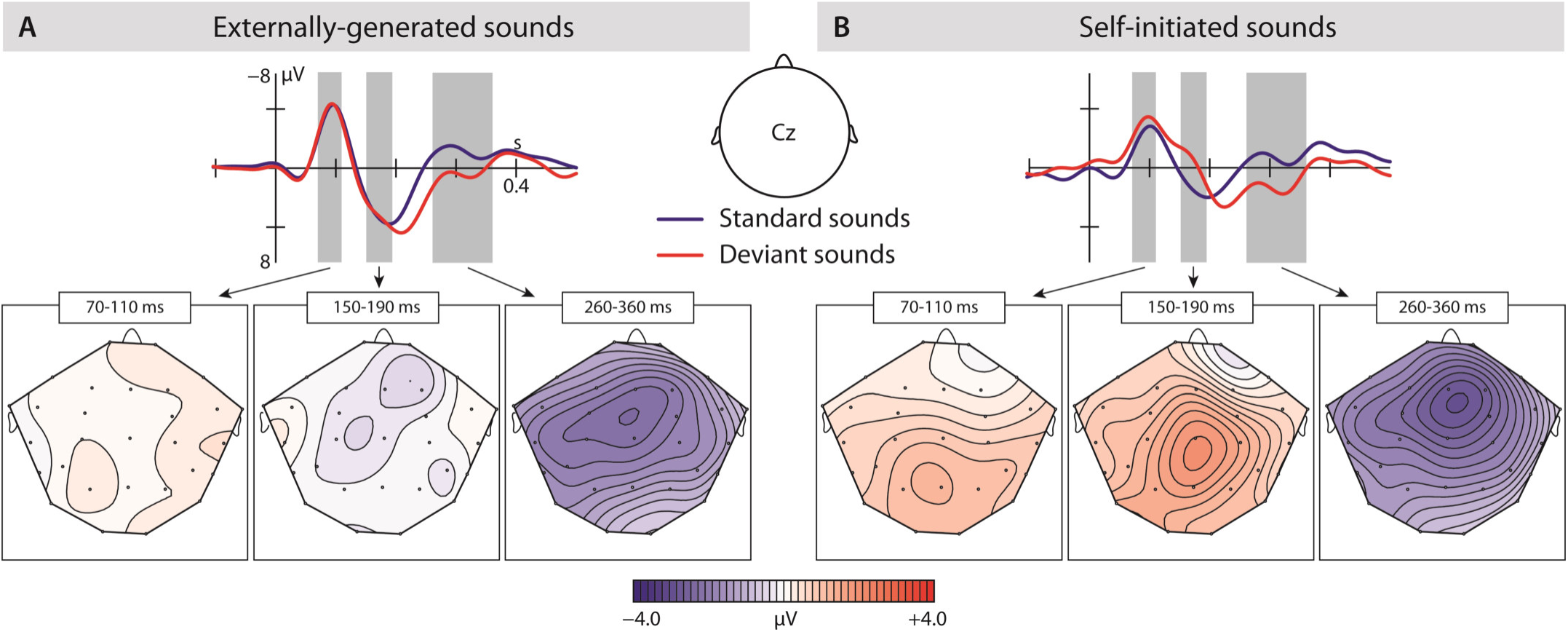
A) Brain responses elicited by externally-produced standard and deviant vowels in the central region. The blue solid line represents externally-produced standard vowels (VOS-2), whereas the red solid line shows responses elicited by externally-produced deviant vowels (VOD-2). B) Brain responses elicited by self-initiated standard and deviant vowels also in the central region. The blue solid line shows self-produced standard vowels (VCS-2), and the red solid line shows responses to self-initiated deviant vowels (VCD-2). The brain maps show the distribution of the effect (for externally-produced vowels: VOS-2 minus VOD-2; for self-initiated vowels: VCS-2 minus VCD-2). Grey bar reflect analyzed in the time window of the N1, N2b and P3a

Taken together, the results show a significant P2-reduction in response to self-initiated standard sounds compared to externally-produced standards. Although externally-produced deviant and standard vowels elicited a similar P2 effect, a possible P2 in response to self-initiated deviant vowels cannot be statistically evaluated due to potential overlay effects.

In the N2b time window we found a significant difference between conditions (self-initiated, externally-generated) and types (standard, deviant), as well as a significant interaction between condition and region, type and region, condition and type, and condition, type and region. The post-hoc analysis of condition, revealed a significant difference between vowel types, as well as a significant interaction between vowel type and region, showing that only self-initiated deviant vowels elicited an N2b effect (standard vowels: VCS-2 mean: -.29 µV; deviant vowels: VCD-2 mean: −1.48 µV).

In contrast (Figure 4a), we did not find a significant difference when comparing externally-produced standard and deviant vowels, indicating similar processing effort, also seen in the mean amplitude values (standard vowel: VOS-2 mean:. 85 µV; deviant vowels: VOD-2 mean:. 69 µV).

In conclusion, the N2b was only elicited in response to self-initiated deviant vowels.

In the P3a time window (Figure 3b; 4 a&b), we found a significant difference between the vowel types (TYPE *F*(1,15) = 22.68; *p* = .0005). In a post-hoc analysis we resolved the type effect, analyzing standard and deviant sounds separately. We found a significant difference between the two conditions in the deviant vowels (CONDITION *F*(1,15) = 5.24; *p* = .037), revealing a significantly enhanced P3a in response to self-initiated deviant vowels (VCC-2 mean amplitude: 1.60 µV) compared to externally-produced deviant vowels (VOC-2 mean amplitude:. 64 µV) (Figure 3b). However, we did not find a significant difference between self-initiated standard vowels and externally-produced standard vowels (VCC-2 mean amplitude: -.44 µV, VOC-2 mean amplitude: -.48 µV) (Figure 3a). The result suggested that only deviant vowel elicited a significant P3a effect which was significantly enhanced in response to self-initiated compared to externally-produced deviant vowels.

## Discussion

The current study investigated the questions whether precise predictions are generated to process self-initiated complex speech sounds (i.e., vowels) by applying an internal forward model and whether prediction errors regarding self-generated deviant vowels are reflected in corresponding ERP components. In order to address these questions, two experiments were conducted. In the first experiment, we used a standard N1-suppression paradigm comparing self- and externally-produced individually pre-recorded vowels in order to test whether the N1-suppression paradigm suffices to investigate complex auditory stimuli such as speech sounds. The results revealed a strong N1-suppression effect in response to self-initiated vowels. This finding confirms the successful generation of a forward prediction independent of the complexity of an anticipated stimulus.

Moreover, we found a reduced P2 response elicited by self-initiated vowels. Although the literature is very diverse regarding the P2, with some studies showing a suppression effect (Houde et al., 2002; Knolle et al., 2012, 2013a and b; Wang et al., 2014), whereas others do not (Baess et al., 2008; Behroozmand, Liu, & Larson, 2011; Chen et al., 2013; Martikainen et al., 2005), the pattern we find in the current data is comparable to our previous results utilizing sounds (Knolle et al., 2012; Knolle et al., 2013a, 2013b). Thus, we suggest that the present data may indicate two processing stages of forming a prediction: Whereas the N1 reflects a fast and automatic forward prediction that prepares the auditory cortex to receive predicted sensory input, the P2 effect represents a more cognitive response (De Chicchis et al., 2002; Crowley & Colrain, 2004), as in distinguishing self-from externally-produced vowels by consciously detecting a self-initiated sensation. This notion is supported by patient studies (Knolle et al., 2012, 2013a). The results of these studies revealed that patients with cerebellar lesions did not show an N1-suppression effect in response to self-initiated sinusoidal sounds, indicating an inability to generate a fast and automatic forward prediction. However, the patients showed a reduced P2 indicating that they had consciously recognized a respective sound as a self-generated one.

Interestingly, a recent study (Pinheiro et al., 2018) testing non-clinical voice hearers using a similar setup as the first experiment in the current study, also using natural vowel sounds, showed that the N1-supression effect is reversed in non-clinical voice hearers with high symptom scores compared to those with low symptom scores whereas the P2-supression effect is maintained in both groups. This is in accordance with findings in schizophrenia patients (Ford, et al., 2014) using a similar paradigm, which indicates that alterations in generating motor-to-auditory predictions might be linked to developing auditory hallucinations (Brebion et al., 2016; Pinheiro et al., 2017).

In the second experiment, we adapted the N1-suppression paradigm by comparing self- and externally-produced standard and deviant vowels. The deviant vowels (30%) were either an /a:/ or an /o:/ dependent on which of these two vowels represented the standard vowel. Comparable to the results of the first experiment, standard vowels elicited a suppressed N1 and P2 component in response the self-initiated vowels indicating that motor-to-auditory predictions are generated in order to process self-initiated vowels. Self-initiated deviant vowels, on the other hand, did not elicit a clear N1-suppression effect, indicating the violation of a precise prediction. In contrast, externally-produced deviants did not differ from externally-produced standard vowels showing that the difference in the N1-suppression effect was not elicited due to deviant detection. Additionally, our results show that the P2 in response to deviant vowels reveals a more complicated pattern: The P2 elicited by externally-produced deviants is well pronounced and very similar to the P2 in response to externally-produced standard vowels. In contrast, the potential P2 component in response to self-initiated deviant vowels is overlaid by an N2b response (Näatänen et al., 1982; for a review, see Näatänen & Gaillard, 1983), and cannot easily be interpreted.

Furthermore, deviant vowels elicited an N2b effect, which suggests the conscious detection of an unexpected, infrequent stimulus (Horvath et al., 2008; Näätänen et al., 1982). The effect was enhanced in response to self-compared to externally-generated deviants. A further study (Kotz, Stockert, & Schwarzte, 2014) revealed an increased N2b response to deviant sounds. Although Kotz and colleagues (2014) did not use a self-generation paradigm they investigated predictability by changing timing and context information to create different degrees of predictability. Their results show that irregular deviants elicit the biggest N2b response showing the detection of a prediction error. This is in accordance to our finding which provides further support for the notion that precise predictions in response to complex stimuli are generated, because only if a prediction concerning a specific feature of the auditory input (i.e., vowel quality) or temporal structure exists, its violation can be detected faster compared to unpredictable auditory input. As these deviants create a prediction error, the result suggests that prediction errors with respect to self-initiated stimuli are more salient, as a self-initiated stimulus is still temporally predictable. This suggestion is supported by our findings of a P3a response to infrequent, unexpected stimuli, to which attention is drawn (Linden, 2005; Polich, 2007; Snyder & Hillyard, 1976; Squires et al., 1975). Here, we report an enhanced P3a effect in response to self-initiated deviants, providing further evidence for the saliency of self-initiated prediction errors (Ford et al., 2010; Nittono & Ullsperger, 2000).

The results of the current study replicated the pattern of components found in a previous study on deviancy processing in sinusoidal sounds (Knolle et al., 2013b). This strongly suggests that predictive processing and the detection of prediction errors occurs independently of stimulus complexity. Additionally, the results reveal that the suppression paradigm is applicable to complex, speech-like sounds, suggesting that the internal forward model provides a theoretical explanation for processing self-produced speech. It can be postulated that by applying an internal forward model, the amplitude of the N1 is modulated when a speech sound is self-initiated compared to when this same speech sound is externally-triggered (Paus et al., 1996; Ventura et al., 2009). In the same line of thought, many studies, using different methods and paradigms, compared spontaneous, self-produced speech to externally-produced speech. They consistently find that spontaneously self-produced speech elicits suppressed cortical responses compared to recorded speech (fMRI: Christoffels, Formisano, & Schiller, 2007; Christoffels et al. 2011; Hashmoto & Sakai, 2003; MEG: Aliu, Houde, & Nagarajan, 2009; Curio, et al. 2000; Heinks-Maldonado, Nagarajan, & Houde, 2006; Houde, Nagarajan, Sekihara, & Merzenich, 2002; Kauramäki, Jääskeläinen, Hari, Möttönen, Rauschecker, & Sams, 2010; Ventura et al., 2009; EEG: Ford, Mathalon, Heinks, Kalba, Faustman, & Roth, 2001; Heinks-Maldonalo, Mathalon, Gray, & Ford, 2005).

However as we hypothesized that the N1-suppression effect reflects the precision of the prediction, we modulated the standard paradigm introducing deviant vowels (second experiment) to reduce the prior knowledge concerning an anticipated stimulus and engenders a prediction error. The results show that precise predictions are generated (Bendixen, SanMiguel & Schröger, 2012; Ford & Mathalon, 2012; Knolle et al., 2013b) as the N1-suppression effect is modified in response to self-initiated deviant vowels. We propose that a prediction generated to process a self-initiated vowel holds a concrete representation of the vowel including for example, its frequency, onset, and intensity. This prediction is violated when a deviant is elicited. It is less precise, as it contains incorrect information on a vowel’s acoustic quality. However, as the concept is still correct with regard to vowel intensity and temporal occurrence, the suppression effect is maintained but modulated. The vocalization literature investigating altered auditory feedback consistently reports a similar effect. For example, Behroozmand and colleagues (2011) reported a reduced N1-suppression effect in response to self-produced but pitch- or onset-altered vocalizations. Thus, participants formed a precise prediction concerning the temporal and acoustic appearance of their voice. When the auditory feedback was altered in frequency or in onset, the prediction was violated. Consequently, the N1-suppression effect was reduced in response to altered auditory feedback, compared to unchanged feedback. Interestingly, two very similar MEG studies show a reduction of the suppression effect of the N1m with regard to altered self-produced speech sounds (Niziolek, Nagaraja, & Houde, 2013; Ylinen et al., 2014),. In accordance to our interpretation Ylinen and colleagues (2014) argue that the reduced suppression reflects a precise motor-to-auditory forward prediction used for concrete speech monitoring. Based on very similar results, Niziolek and colleagues (2013) argue that specific speech monitoring allows error detection as well as concrete error correction mechanisms, which is in accordance to our interpretation.

The assumption that precise predictions are generated also in response to complex self-initiated stimuli receives further support from the enhanced N2b effect in response to self-initiated deviant vowels. Similar to our previous study (Knolle et al., 2013b), we consider that the N2b indicates the conscious detection of a prediction error, which represents an infrequent stimulus (Horvath et al., 2008; Näätänen et al., 1982). As the N2b is very much enhanced in response to self-initiated deviant vowels, it can be suggested that a prediction error increased the saliency of a deviant (Ford et al., 2010). More generally speaking, when a prediction error is generated, the detection of such violation is processed more efficiently in self-produced compared to externally-produced deviant vowels. This is supported by our finding that externally-produced deviant vowels also show a reduced P3a compared to self-produced deviants, revealing a less salient response (Ford et al., 2010).

The results concerning the detection of deviants nicely complements the results presented in the literature on speech monitoring and processing of speech errors (Christoffels et al., 2011; Postma, 2000; Tourville, Reilly, & Guenther, 2008; Zheng, Munhall, & Johnsrude, 2010) which most reliably show an increased blood oxygenation level dependent (BOLD) response during altered auditory feedback compared to normal feedback, indicating that increased activity is necessary to accomplish the monitoring effort or error coding. In the current study we show that violated predictions – prediction errors – are reflected in specific ERP components (i.e., enhanced N2b and P3a), which can be compared to error coding in fMRI studies.

In the current study we have investigated motor-to-auditory predictions, framed within a forward model account, which is specific to motor control and uses the concept of a efference copy to generate a prediction. Other current accounts of surprising (e.g., oddball or deviant) responses in a more general, modality-independent framework are usually cast in terms of predictive coding (e.g. Brown et al. 2013; Friston, 2018; Shipp, 2016). In these models, the brain uses a hierarchical forward or generative model inferring the causes of sensations from its sensory consequences. Violations or mismatches from these predicted sensations result in a prediction error that can be differently weighted based on their precision (Haarsma et al., 2018) and is used for belief updating in cortical hierarchies. How does this concept link to agency? To explain self-produced action, such as speech, current accounts treat motor commands as predictions of proprioceptive and somatosensory consequences of an intended act, while the efference copy or corollary discharge corresponds to the predictions in the exteroceptive (e.g. auditory or visual) domain (Sterzer et al., 2018). Crucially, to act, it is necessary to attenuate the gain or precision of ascending prediction errors. Psychologically, this manifests as sensory attenuation as measured psychophysically. Physiologically, this is usually manifest as an attenuation or suppression of evoked responses that are generated by self, relative to another, as shown in this study.

In conclusion, the present study presents two experiments: The first investigates whether forward predictions are generated to process self-initiated complex speech sounds (i.e., vowel)? And the second addresses the question whether prediction error regarding self-initiated deviant, are these predictions based on precise patterns, as in holding a concrete representation of the anticipated vowel? Addressing the first question, the results revealed N1 and P2 suppression elicited by self-initiated vowels, indicating a successful generation of predictions in response to complex self-initiated speech sounds. This finding supports the notion that processing of self-produced speech mirrors components of a forward model, preparing respective cortical areas for incoming sensory consequences of self-produced speech. Investigating the second question, we found N1-suppression in response to self-initiated vowels compared to externally-generated vowels. In addition, we report an enhanced N2b and P3 effect in response to self-initiated compared to externally-produced deviant vowels. These findings imply that specific predictions are generated. Furthermore, we show that prediction errors are more salient in self-initiated speech sounds compared to externally-produced sounds. More generally, our results speak to a key role of agency in predictive processing formulations; namely, a key role in mediating sensory attention and its reversal when attending to the consequences of self-generated acts.

## Acknowledgement

The research was funded by a DFG-Reinhart-Koselleck grant to ES and a DFG KO 2268/6-1 grant to SAK. We thank K. Ina Koch and Eleni Beyer for their support in data collection. Furthermore, we thank Kerstin Flake for graphics support and Helga Smallwood for proofreading.

